# Phenolic glycolipid facilitates mycobacterial escape from a microbicidal population of tissue-resident macrophages

**DOI:** 10.1101/147421

**Authors:** C.J. Cambier, Seónadh M. O’Leary, Mary P. O’Sullivan, Joseph Keane, Lalita Ramakrishnan

**Affiliations:** Department of Immunology, University of Washington, Seattle, USA; Department of Medicine, University of Cambridge, Cambridge, UK; Department of Chemistry, Stanford University, Stanford 94305, USA; Department of Clinical Medicine, Trinity Translational Medicine Institute, Trinity College Dublin, Dublin 8, Ireland; Department of Microbiology, University of Washington, Seattle, USA; Department of Medicine, University of Washington, Seattle, USA

## Abstract

*Mycobacterium tuberculosis* enters the host in aerosol droplets deposited in lung alveoli where the bacteria first encounter lung-resident alveolar macrophages. We studied the earliest mycobacterium-macrophage interactions in the optically transparent zebrafish. We find that the first-responding resident macrophages can phagocytose and eradicate infecting mycobacteria. So, to establish a successful infection, mycobacteria must escape out of the initial resident macrophage into growth-permissive monocytes. We define a critical role for the membrane phenolic glycolipid (PGL) in engineering this transition to a permissive niche. PGL activates the STING cytosolic sensing pathway, thereby inducing the chemokine CCL2 that recruits permissive peripheral monocytes. The bacteria then transfer from resident macrophage to recruited monocyte via transient fusion of the two immune cells. We show that interrupting this bacterial strategy so as to prolong the mycobacterial sojourn in resident macrophages promotes clearing of infection. Because PGL-dependent CCL2 induction is conserved in human alveolar macrophages, our findings suggest the potential of immunological or pharmacological PGL-blocking interventions to prevent tuberculosis.

## INTRODUCTION

When *M. tuberculosis* is aerosolized into the lower lung, it first encounters lung resident alveolar macrophages that patrol the air - lung epithelium interface (Srivastava et al., 2014). *M. tuberculosis* is found for the first few days exclusively within alveolar macrophages (Srivastava et al., 2014; Urdahl, 2014; Wolf et al., 2007). Thereafter, it is found to have traversed the lung epithelium to be within other myeloid cells that have aggregated into granulomas (Cambier et al., 2014a; Srivastava et al., 2014). The difficulty of tracking the early fate of individual mycobacteria in traditional animal models has precluded elucidation of how mycobacteria move from alveolar macrophages into other cells, and indeed how they survive these broadly microbicidal first-responders (Hocking and Golde, 1979).

We have exploited the optical transparency of the zebrafish larva to study the early mycobacterium-phagocyte interactions by infecting *Mycobacterium marinum*, a close genetic relative of *Mycobacterium tuberculosis*, into the zebrafish larval hindbrain ventricle, an epithelium-lined cavity (Fig. 1a) (Cambier et al., 2014b; Yang et al., 2012). We have shown previously that immediately upon infection, pathogenic mycobacteria manipulate host responses so as to inhibit the recruitment of neutrophils and microbicidal monocytes, and instead to recruit and infect mycobacterium-permissive myeloid cells (Cambier et al., 2014b; Yang et al., 2012). We have identified the strategy by which pathogenic mycobacteria avoid detection of microbicidal monocytes: they mask their bacterial pathogen associated molecular patterns (PAMPs) with the cell-surface phthiocerol dimycoceroserate (PDIM) lipid so as to prevent detection by toll-like receptors (TLRs) (Cambier et al., 2014b). They thus inhibit monocyte signaling through TLRs, which recruits prototypical microbicidal iNOS-expressing monocytes.

**Figure 1:**
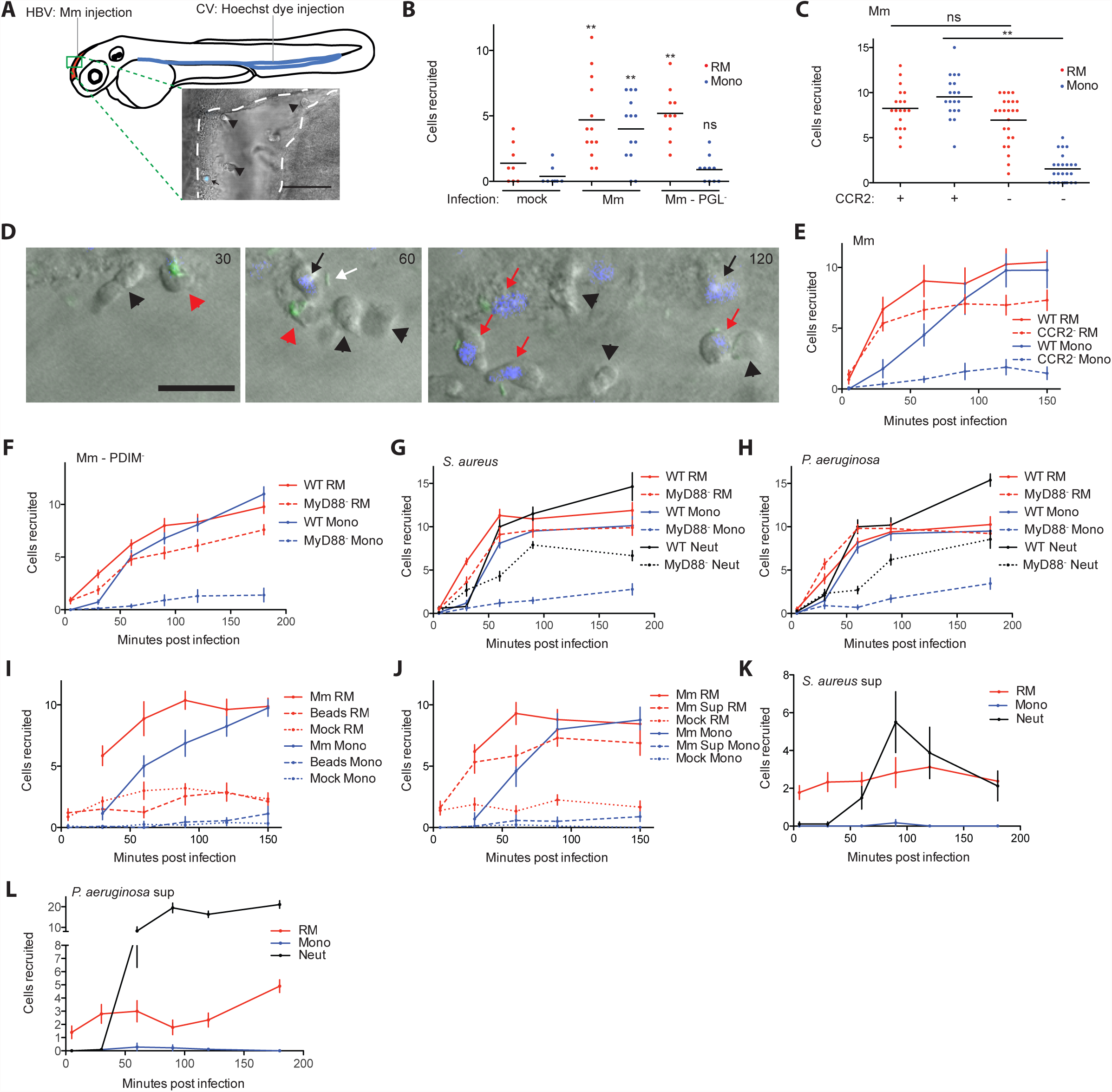
Resident macrophages are first responders to bacterial infection. (A) Cartoon of a 2 day post-fertilization (dpf) zebrafish showing the caudal vein (CV) and hindbrain ventricle (HBV) injection sites. (B) Mean resident macrophage (RM) and monocyte (Mono) recruitment at 3hours post infection (hpi) into the HBV after infection with 80 wildtype *M. marinum* (Mm) or PGL-deficient *M. marinum* (Mm-PGL^-^). Significance testing done using one-way ANOVA, with Bonferroni’s post-test against mock injections. \*\**P* < 0.01. (C) Mean resident macrophage and monocyte recruitment at 3hpi into the HBV of wildtype or CCR2-deficient fish after infection with 80 wildtype *M. marinum.* Significance testing done using one-way ANOVA, with Bonferroni’s post-test for comparisons shown. \*\**P* < 0.01. (D) Representative images of uninfected resident macrophages (black arrowheads), uninfected monocytes (black arrows), infected resident macrophages (red arrowheads), infected monocytes (red arrows), and extracellular bacteria (white arrow) following infection of wildtype fish in the HBV with 80 wildtype green fluorescent *M. marinum* at 30, 60, and 120 minutes post infection (mpi). Scale bar, 20μm. (E) Mean resident macrophage and monocyte recruitment from 5 to 150mpi in the HBV of wildtype or CCR2-deficient fish after infection with 80 wildtype *M. marinum*. (F) Mean resident macrophage and monocyte recruitment from 5 to 180mpi in the HBV of wildtype or MyD88-deficient fish after infection with 80 PDIM-deficient (PDIM^-^) *M. marinum*. (G and H) Mean resident macrophage, monocyte, and neutrophil (Neut) recruitment from 5 to 180 mpi in the HBV of wildtype or MyD88-deficient fish following infection with 138 *S. aureus* (G) or 156 *P. aeruginosa* (H). (I) Mean resident macrophage and monocyte recruitment from 5 to 150mpi in the HBV of wildtype fish after injection with 80 wildtype *M. marinum*, 300 sterile beads, or mock injection. (J) Mean resident macrophage and monocyte recruitment from 5 to 150mpi in the HBV of wildtype fish after infection with 80 wildtype *M. marinum*, an equivalent volume of wildtype *M. marinum* supernatant (Sup), or media mock. (K and L) Mean resident macrophage, monocyte, and neutrophil recruitment from 5 to 180mpi in the HBV of wildtype fish after infection with *S. aureus* supernatant (K) or *P. aeruginosa* supernatant (L). (A – L) Representative of at least three separate experiments.

Here, we find that these elaborate evasion strategies notwithstanding, mycobacteria still have to contend with first-responding resident macrophages that they cannot avoid. We show that resident macrophages are default first responders to invading bacteria, and pathogenic mycobacteria are no exceptions. These first-responding resident macrophages are microbicidal to virulent mycobacteria and are capable of eradicating infection unless the mycobacteria can escape out of them rapidly into more permissive cells.We show that mycobacteria use a PDIM-related surface lipid, phenolic glycolipid (PGL), to rapidly induce the monocyte chemokine CCL2 in the resident macrophages they infect through STING activation in them. Mycobacterium-permissive monocytes are recruited through signaling by the CCL2 cognate receptor CCR2. The bacteria then transfer into these monocytes, which provide them a growth-permissive niche to establish infection. Our findings identify the resident macrophage-mycobacterium interaction as possibly the earliest determinant of whether infection will be established or cleared, and it reveals PGL as a very early mycobacterial immune evasion determinant. PGL’s partnership with STING and CCL2 for this immune evasion strategy reveals them to be host susceptibility factors that act at the very first steps of infection.

## RESULTS

### Resident Macrophages are First Responders to *M. marinum* and Mucosal Commensal Pathogens Through Sensing a Common Secreted Signal

When *M. tuberculosis* is aerosolized into mouse lung, it is found for the first few days exclusively within alveolar macrophages (Srivastava et al., 2014; Urdahl, 2014; Wolf et al., 2007). In the zebrafish larva, directly posterior to the hindbrain ventricle infection site (Figure 1A), is the brain which, like most organs, has a population of resident macrophages (Herbomel et al., 2001). We asked if these brain-resident macrophages or microglia, analogous to the resident macrophages of the mammalian lung, participated in the immune response toward mycobacterial infection. In addition to their tissue-specific functions, tissue-resident macrophages, including those of the brain, play a central role in host defense against infection (Casano and Peri, 2015). Like lung-resident macrophages, brain-resident macrophages phagocytose *M. tuberculosis* and produce inflammatory cytokines in response to it (Curto et al., 2004; Spanos et al., 2015).

To distinguish between brain-resident macrophages and monocytes, we took advantage of the nuclear dye Hoechst 33342 that does not cross the blood brain barrier, so that its injection into the caudal vein of zebrafish larvae labels cells, including myeloid cells, in the body but not in the brain (Davis and Ramakrishnan, 2009). We injected Hoechst dye into the caudal vein, then two hours later *M. marinum* into the HBV (Figure 1A). Three hours following infection, recruited cells were identified as either brain-resident macrophages (Hoechst-negative) or peripheral monocytes (Hoechst-positive) (Figure 1A). Wildtype *M. marinum* recruited both resident macrophages and monocytes whereas the PGL-deficient *M. marinum* strain recruited resident macrophages but not monocytes (Figure 1B). Correspondingly in CCR2-deficient animals, wildtype *M. marinum* recruited resident macrophages but not monocytes (Figure 1C). We asked if resident macrophages in the zebrafish larvae arrived more rapidly to mycobacteria, similar to resident macrophages in the mammalian lung. A temporal analysis revealed that they were the first responders to infection and arrived independently of CCR2 signaling (Figure 1D and 1E). In contrast, monocytes arrived later and in a CCR2-dependent fashion (Figure 1D and 1E). Thus, similar to *M. tuberculosis* infection of the mammalian lung, *M. marinum* infection of the zebrafish HBV recruits both resident macrophages and peripheral monocytes. The two cell types appear to be recruited sequentially, and through distinct pathways - CCR2-independent for resident macrophages and CCR2-dependent for peripheral monocytes.

We found that resident macrophages were also the first-responders to bacteria to which overall myeloid cell recruitment is dependent on toll-like receptor (TLR/MyD88) signaling rather than CCL2/CCR2 (Cambier et al., 2014b) - e.g. PDIM-deficient *M. marinum* (Δ*mmpL7*) and the mucosal commensal-pathogens *Staphylococcus aureus* and *Pseudomonas aeruginosa* (Figure 1F-1H). *S. aureus* and *P. aeruginosa* are known to elicit early recruitment of neutrophils in addition to mononuclear phagocytes, both through TLR/MyD88 signaling (Figure 1G and 1H) (Cambier et al., 2014b; Yang et al., 2012). In all cases, resident macrophage recruitment was independent of TLR/MyD88 signaling (Figure 1F-1H). Taken together, our data suggested that tissue resident macrophages are default first responders to invading bacteria, even those that elicit a robust protective neutrophilic response. Furthermore, they are signaled by TLR-independent pathways. We ruled out the possibility that mechanosensing of a foreign body at the infection site was driving resident macrophage recruitment (Wang et al., 2009) by showing that resident macrophages were recruited towards sterile beads (and neither were monocytes) (Figure 1I). These data suggested that resident macrophage recruitment is specifically mediated by bacterial signals. We then showed that supernatants of *M. marinum*, *S. aureus* and *P. aeruginosa* all recruited resident macrophages (and in the case of the latter two, neutrophils) but not monocytes (Figure 1J-L). Thus, tissue-resident macrophages are default first-responders signaled by a pathway that is both TLR/MyD88- and CCL2-independent, in response to a secreted factor(s) produced by a variety of bacteria independent of their Gram reaction.

### Mycobacteria actively induce CCL2 in resident macrophages to recruit CCR2- dependent monocytes

For mycobacterial infection, our findings that resident macrophages are rapidly recruited through a PGL/CCL2-independent pathway followed by PGL/CCL2-dependent monocyte recruitment, led us to ask if monocyte recruitment was dependent on resident macrophage recruitment. We used zebrafish larvae depleted of myeloid cells (by morpholino knockdown of the myeloid transcription factor PU. 1 (Clay et al., 2007)) and evaluated *ccl2* RNA expression following infection with PGL-expressing *M. marinum.* Myeloid-deficient fish were unable to induce *ccl2* (Figure 2A). This finding suggested that resident macrophages are required for CCL2 expression. Next, we asked if *ccl2* expression was resident macrophage-intrinsic, using in-situ hybridization analysis with a *ccl2* antisense RNA probe (Clay et al., 2007). We found that the macrophages themselves express *ccl2*, and specifically in response to PGL-expressing bacteria: at one hour post infection, when the recruited phagocytes comprise mostly resident macrophages (Figure 1E), *ccl2*-positive phagocytes were present, but only following WILDTYPE *M. marinum* infection and not PGL-deficient *M. marinum* infection (Figure 2B-2D). Next, to directly test if resident macrophages are required for monocyte recruitment, we used zebrafish mutants in which colonization of the brain by resident macrophages is delayed due to a genetic mutation in colony stimulating factor receptor 1 (CSF1R) (Herbomel et al., 2001). Therefore, at the time of our recruitment assay (two days post fertilization), *csf1r*^-/-^ fish have normal numbers of circulating monocytes but very few resident macrophages (Herbomel et al., 2001; Pagán et al., 2015) (Figure 2E). Following wildtype *M. marinum* infection into the HBV of *csf1r*^-/-^ fish, resident macrophage recruitment was decreased and delayed, consistent with the lack of available cells in the brain (Figure 2F). Importantly, monocyte recruitment was also markedly decreased, consistent with our hypothesis that resident macrophages mediate monocyte recruitment (Figure 2F). In conjunction with our earlier finding that mycobacterial PGL was also required for monocyte recruitment (Figure 1B), these findings supported a model where resident macrophages, recruited in response to generic bacterial signals, engulf the mycobacteria. Mycobacterial PGL then induces them to express CCL2 that mediates monocyte recruitment. Because PGL is heat-stable (Onwueme et al., 2005), this model would predict that heat-killed PGL-expressing *M. marinum* would both induce CCL2 and recruit monocytes. It did neither, suggesting that live PGL-expressing mycobacteria are required to recruit monocytes through CCL2 induction in resident macrophages (Figure 2G and 2H). Notably, heat-killed bacteria did in fact recruit resident macrophages (Figure 2H) consistent with the secreted factor responsible for resident macrophage recruitment being heat-stable.

**Figure 2:**
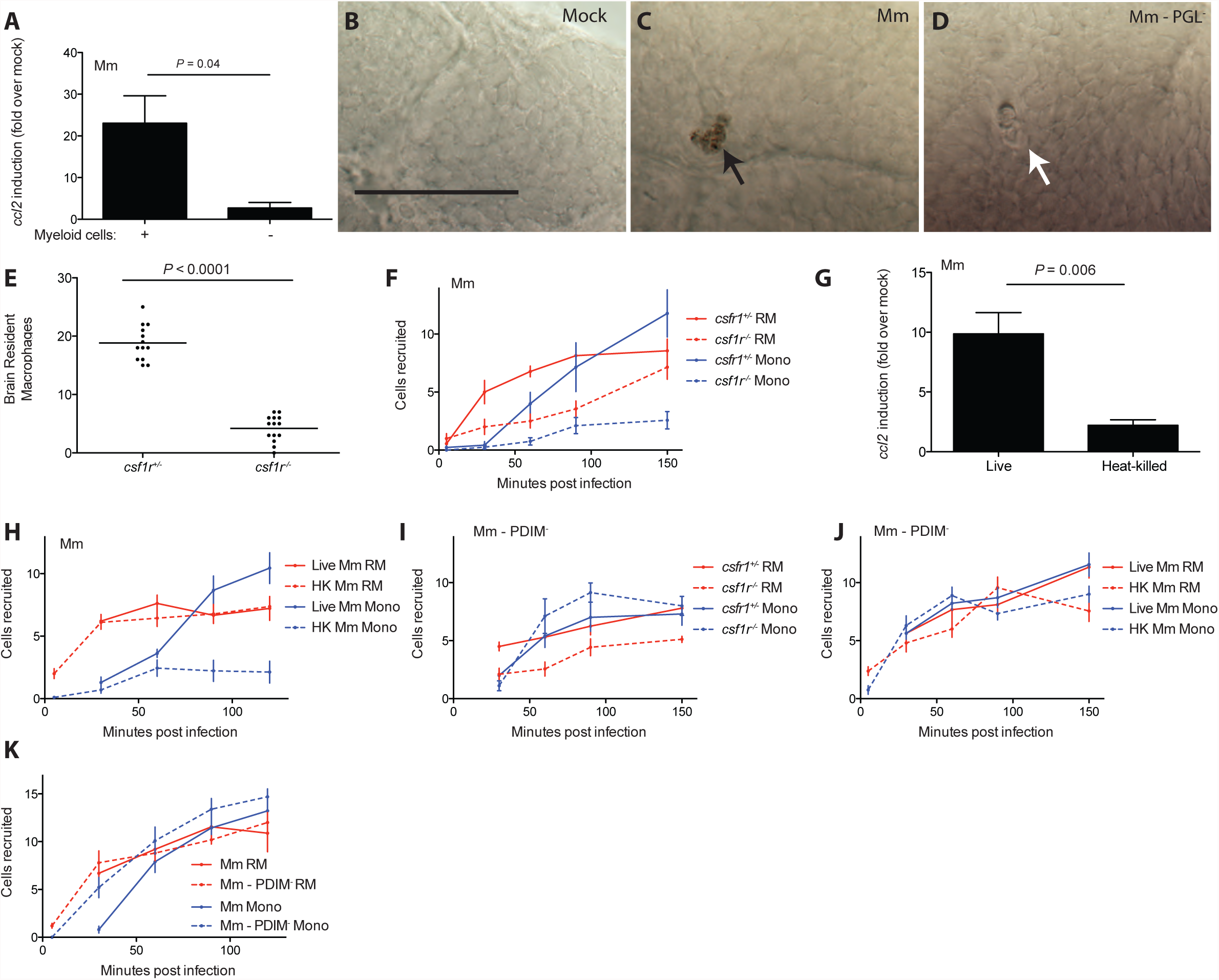
Mycobacteria mediate CCR2-dependent monocyte recruitment by actively inducing CCL2 in resident macrophages. (A) *ccl2* messenger RNA levels (mean +/- s.e.m. of three biological replicates) induced at 3 h after caudal vein infection of 2 dpf wildtype or myeloid cell-deficient fish with 250–300 wildtype wildtype *M. marinum*. (B-D) In situ hybridizations against zebrafish *ccl2* mRNA following hindbrain ventricle infections with vehicle (bacterial media) (B), 80 wildtype *M. marinum* (C), 80 PGL^-^ *M. marinum* (D). Black arrows, *ccl2* mRNA-positive phagocytes; white arrows *ccl2* mRNA-negative phagocytes. (E) Mean brain resident macrophage numbers of *csfr1*^+/-^ and *csfr1*^-/-^ zebrafish at 2dpf. (F) Mean resident macrophage and monocyte recruitment from 5 to 150 mpi in the HBV of *csfr1*^+/-^ or *csfr1*^-/-^ fish after infection with 80 wildtype *M. marinum*. (G) *ccl2* messenger RNA levels (mean +/- s.e.m. of three biological replicates) induced at 3 h after caudal vein infection of 2 dpf wildtype fish with 250–300 live or heat-killed wildtype *M. marinum*. (H) Mean resident macrophage and monocyte recruitment from 5 to 120mpi in the HBV of wildtype fish after infection with 80 live or heat-killed (HK) wildtype *M. marinum*. (I) Mean resident macrophage and monocyte recruitment from 5 to 150mpi in the HBV of *csfr1*^+/-^ or *csfr1*^-/-^ fish after infection with 80 PDIM- *M. marinum*. (J) Mean resident macrophage and monocyte recruitment from 5 to 150mpi in the HBV of wildtype fish after infection with 80 live or heat-killed (HK) PDIM- *M. marinum* (K) Mean resident macrophage and monocyte recruitment from 5 to 120mpi in the HBV of wild-type fish after infection with 80 wildtype or PDIM- *M. marinum*.

Our finding that peripheral monocytes were dependent on signals from resident macrophages to participate in mycobacterial infection was surprising, and we wondered if this requirement was unique to PGL-expressing mycobacteria. To test this, we used PDIM-deficient *M. marinum*, which recruits monocytes through TLR/MyD88, not CCL2. *Csf1r*^-/-^ zebrafish recruited monocytes normally to PDIM-deficient bacteria (Figure 2I). Moreover, in contrast to wildtype mycobacteria, heat-killed PDIM-deficient *M. marinum* recruited monocytes (Figure 2J). These results suggested a passive detection of the surface-exposed TLR ligands of this mutant bacterium (Cambier et al., 2014b) in contrast to an active recruitment process mediated through live PGL-expressing bacteria. A head-to-head comparison of the recruitment kinetics of wildtype and PDIM-deficient strains revealed earlier monocyte recruitment to PDIM-deficient bacteria (Figure 2K), consistent with their recruitment to this strain being independent of resident macrophages. In sum, resident macrophages specifically promote CCL2-dependent monocyte recruitment in response to virulent mycobacteria, and this is dependent on mycobacterial PGL.

Taken together, our findings suggest that heat-stable bacterial PAMPs of PDIM-deficient *M. marinum* trigger a program of microbicidal monocyte recruitment that is not dependent on resident macrophages. In contrast, when bacterial PAMPs are masked by PDIM, PGL-mediated recruitment of permissive monocytes is absolutely dependent on both resident macrophages and live bacteria, suggesting an active bacterial manipulation of these default first-responders.

### *M. marinum* PGL recruits monocytes through STING-dependent CCL2 induction

How might PGL induce CCL2 in resident macrophages? Because PGL operated in the context of live bacteria, we wondered if a cytosolic sensing pathway was involved. Activation of the cytosolic signaling pathway STING can induce CCL2 (Chen et al., 2011), so we tested if STING was the intermediary in PGL-mediated CCL2 induction. STING depletion using a splice-blocking morpholino (Ge et al., 2015) resulted in a lack of *ccl2* induction in the resident macrophages in response to *M. marinum* (Figure 3A – 3C). Consistent with the inability to induce *ccl2*, STING-deficient animals had reduced monocyte recruitment to *M. marinum* while leaving resident macrophage recruitment intact (Figure 3D). Importantly, STING-deficient animals recruited monocytes normally to PDIM-deficient *M. marinum* confirming that their inability to elicit monocytes was specifically in the context of CCL2-mediated and not MyD88-dependent monocyte recruitment (Figure 3E). Finally, our model would predict that like CCL2/CCR2 deficiency, STING deficiency should compromise the ability of wildtype bacteria to establish infection. Mycobacterial infectivity can be stringently tested by infecting animals with very low inocula that resemble human infection; in the zebrafish we have developed an infectivity assay which determines how many animals remain infected 4-5 days after infection with 1-3 mycobacteria (Cambier et al., 2014b). Using this infectivity assay, we found that wildtype *M. marinum* had reduced infectivity in STING-deficient animals (Figure 3F), similar to PGL-deficient bacteria in wildtype animals and wildtype bacteria in CCR2-deficient animals (Cambier et al., 2014b).

**Figure 3:**
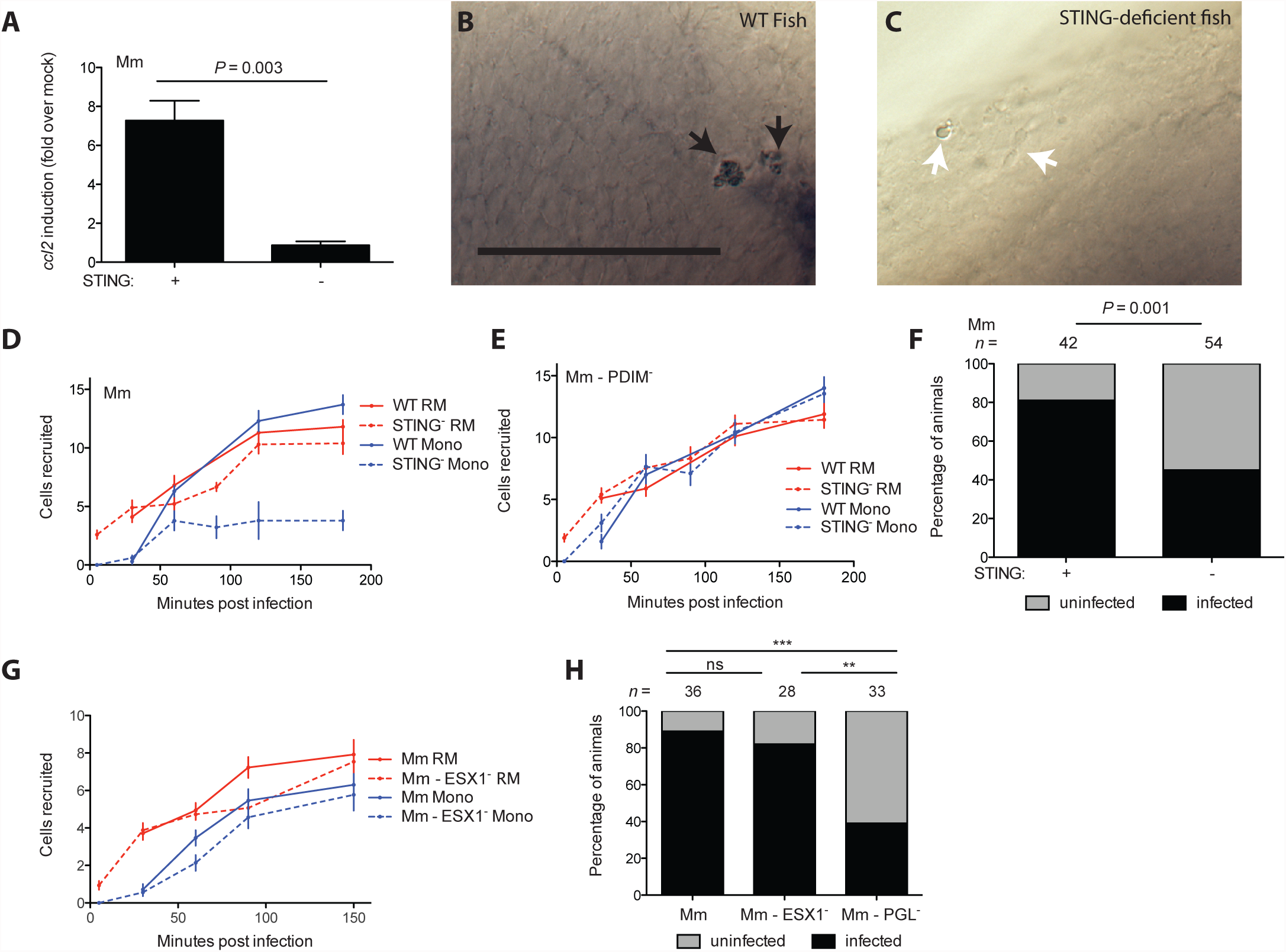
*M. marinum* PGL recruits monocytes through STING-dependent CCL2 induction. (A) *ccl2* messenger RNA levels (mean +/- s.e.m. of three biological replicates) induced at 3 h after caudal vein infection of 2 dpf wildtype or STING-deficient fish with 250–300 wildtype *M. marinum*. Student’s unpaired t-test. (B and C) In situ hybridizations against zebrafish *ccl2* mRNA following hindbrain ventricle infections with 80 wildtype *M. marinum* into wildtype (B) or STING-deficient (C) zebrafish. Black arrows, *ccl2* mRNA-positive phagocytes; white arrows *ccl2* mRNA-negative phagocytes. (D) Mean resident macrophage and monocyte recruitment from 5 to 180 mpi in the HBV of wildtype or STING-deficient fish after infection with 80 wildtype *M. marinum*. (E) Mean resident macrophage and monocyte recruitment from 5 to 180 mpi in the HBV of wildtype or STING-deficient fish after infection with 80 PDIM^-^ *M. marinum*. (F) Percentage of infected (black) or uninfected (gray) wildtype or STING-deficient fish 5 dpi with 1-3 wildtype *M. marinum* into the HBV. *n* = number of larvae per group. Representative of two separate experiments. Fisher’s exact test. (G) Mean resident macrophage and monocyte recruitment from 5 to 150mpi in the HBV of wildtype fish after infection with 80 wildtype or ESX-1-deficient (ESX1^-^) *M. marinum*. (H) Percentage of infected (black) or uninfected (gray) wildtype fish 5 dpi of 1-3 wildtype, ESX1^-^, or PGL^-^ *M. marinum* into the HBV. *n* = number of larvae per group. Significance testing done using Fisher’s exact test for comparisons shown. \*\**P* < 0.01, \*\*\**P* < 0.001. Representative of two separate experiments.Results in (D), (E), (G) representative of three separate experiments.

STING can induce CCL2 either through Type 1 interferons (IFN) (Cepok et al., 2009),(Conrady et al., 2013), or independently of them (Chen et al., 2011). In the context of mycobacterial infection, the ESX-1 secretion system is required to induce the type I IFN, IFNβ and this activation occurs through ESX- 1 dependent rupture of the phagosome (Siméone et al., 2015) that activates STING (Manzanillo et al., 2012). If STING is activating CCL2 through IFNβ, then monocyte recruitment should be ESX-1-dependent. On the contrary, we found that it was not. ESX- 1 mutant bacteria recruited both resident macrophages and monocytes normally to the initially infecting bacteria (Figure 3G). Consistent with this finding, ESX-1-deficient *M. marinum* established infection at wildtype levels (Figure 3H). Our prior work has found that ESX-1 partners with host MMP9 to accelerate macrophage recruitment to the forming granuloma (Volkman et al., 2004). These new findings show initial macrophage recruitment occurs through a distinct mechanism – PGL activation of STING that directly induces CCL2. It is not surprising that this process is ESX-1 independent because of the timing of CCL2 induction (prior to 3 hours post infection) versus ESX- 1 induced phagosome permeabilization which takes ~ 24 hours (Siméone et al., 2015). Given that STING activation can occur downstream of foreign membrane fusion (Holm et al., 2012), our findings may implicate mycobacterial derived membrane vesicles as a potential mechanism underlying PGL-dependent STING activation. Formation of these vesicles requires bacterial viability (Athman et al., 2015) but not ESX-1 (Bhatnagar and Schorey, 2007), and could result in STING activation at these early stages before phagosomal permeabilization enables a different pathway of STING activation. This scenario would fit with the requirement of PGL, a surface lipid in the vesicle for STING activation.

### PGL-expressing bacteria can transfer from resident macrophages to monocytes

Human TB is thought to result from infection with only 1-3 bacteria (Bates et al., 1965; Cambier et al., 2014a; Wells et al., 1948). In the zebrafish, 1-3 *M. marinum* are sufficient to establish infection in the majority of zebrafish larvae provided that bacterial PGL and host STING and CCL2/CCR2 are present; without these factors, infectivity is reduced (Figure 3F) (Cambier et al., 2014b). Therefore, it was important to examine myeloid cell recruitment in response to these low inocula where the role of PGL and CCR2 is most relevant. To enable a detailed temporal analysis of the HBV by time-lapse confocal microscopy, we used *mpeg::yfp* or *mpeg::tdtomato* transgenic zebrafish with fluorescent myeloid cells, again using Hoechst dye to distinguish monocytes from resident macrophages. Imaging each animal every 10 minutes from 1-11 hours post infection, we observed that resident macrophages arrived early whereas monocytes were rarely seen during this period (Figure 4A and Table S1). In contrast, even with these low inocula, both cell types were recruited early to PDIM-deficient mutants (Figure 4A). Accordingly, when we analyzed the phagocytosis event for each bacterium, we found that wildtype bacteria were phagocytosed only by resident macrophages whereas PDIM-deficient bacteria were phagocytosed by both resident macrophages and monocytes (Figure 4B).

**Figure 4:**
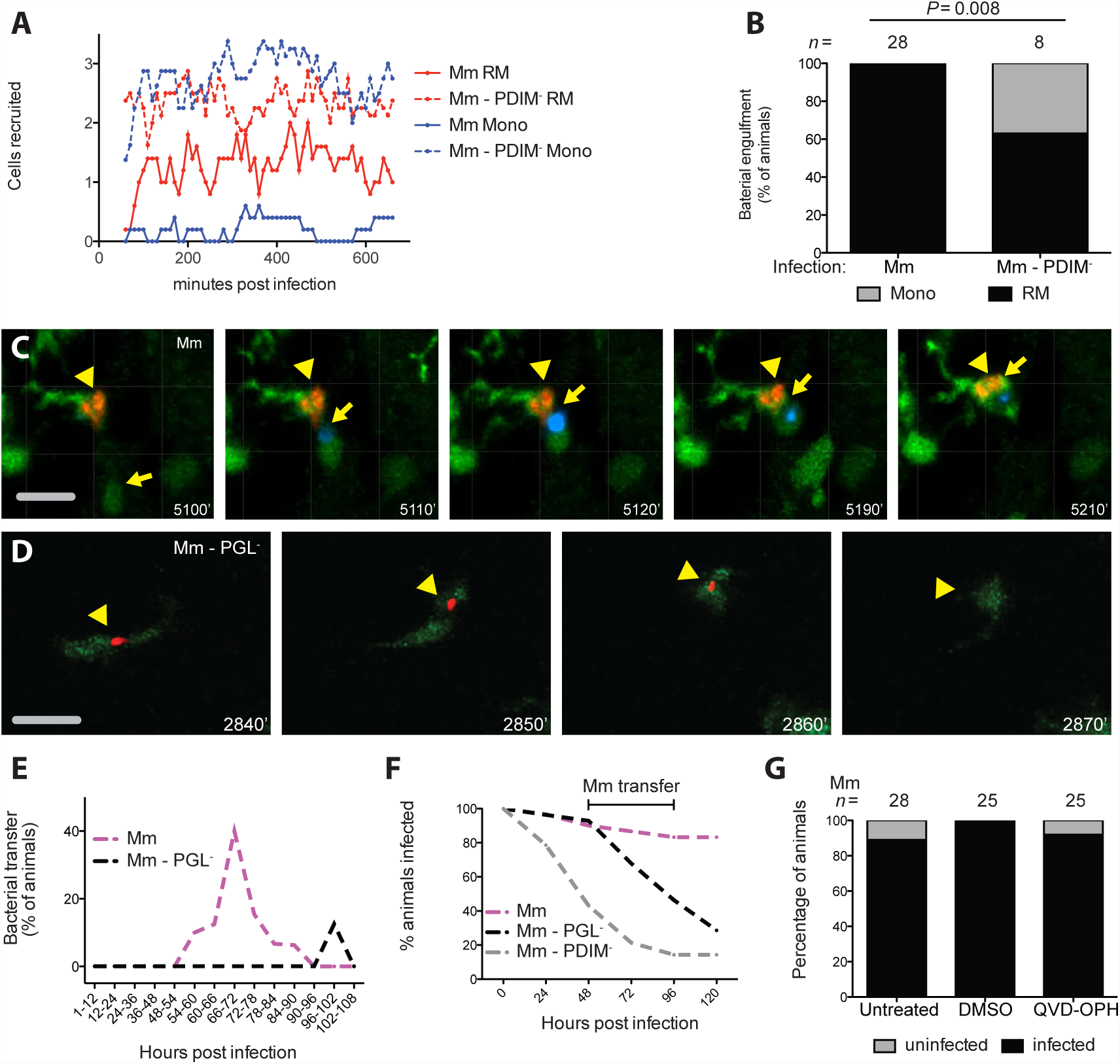
PGL promotes intercellular bacterial transfer and prevents bacterial clearance. (A) Mean (≥5 biological replicates) resident macrophage and monocyte recruitment quantified every 10 min from 1 to 11hpi in the HBV of Tg(*mpeg1::yfp*) fish with green fluorescent macrophages after infection with 1-3 wildtype or PDIM^-^ red fluorescent *M. marinum*. (B) Percentage of fish where the infecting bacteria were phagocytosed by a resident macrophage (black) or a monocyte (gray) over the first 11 hours following infection of Tg(*mpeg1*:YFP) fish in the HBV with red fluorescent 1-3 wildtype or PDIM^-^ *M. marinum*. *n* = number of larvae per group. Fisher’s exact test. (C) Representative images from a time-lapse movie of a bacterial transfer event. Uninfected Hoechst positive (blue fluorescence) monocyte (yellow arrow) is seen phagocytosing an infected cell (yellow arrowhead). Scale bar, 50μm. Time stamp, mpi. (D) Representative images from a time-lapse movie showing an infected macrophage (green fluorescent) clearing red fluorescent PGL^-^ *M. marinum* (yellow arrowhead). Scale bar, 50μm. Time stamp, mpi. (E) Quantification of bacterial transfer events from experiments represented by (C) and (D). Percentage of animals demonstrating a transfer event during the designated imaging time block. (F) Percentage of animals remaining infected over the first 5 days of infection with 1-3 wildtype, PGL^-^, or PDIM^-^ *M. marinum* into the HBV of wildtype fish. Numbers of fish infected with each *M. marinum* strain: 30 wildtype, 28 PGL^-^, and 28 PDIM^-^ *M. marinum*. (G) Percentage of infected (black) or uninfected (gray) untreated, DMSO control, or QVD-OPH treated wildtype fish 5 dpi with 1-3 wildtype *M. marinum* into the HBV. *n* = number of larvae per group. Representative of two separate experiments.

Previously, we had shown that the increased infectivity of PGL-competent bacteria is abrogated by CCR2 deficiency (Cambier et al., 2014b). Now we had found that both PGL-competent and PGL-deficient bacteria are initially in resident macrophages that are recruited in a CCR2-independent manner, with the critical difference between the two strains being whether there is subsequent recruitment of CCR2-dependent monocytes or not. Taken together, the two findings suggested that these monocytes were responsible for the increased infectivity of PGL-competent bacteria. This could be because the monocytes comprised a more permissive niche into which the bacteria were transferring, or because their presence was modulating the microbicidal capacity of the originally-infected resident macrophages.

In order to determine if bacteria were being transferred to new cells, we had to image infection for the first several days. Continuous imaging of the infection site in the same animal for several days is precluded by photobleaching. So we devised a strategy where we divided the infected larvae into 14 groups, and imaged each group for one of consecutive 6-12 hour periods that together spanned 4.5 days of infection (Table S2). For wildtype bacteria, transfer events were observed starting at 54 hours and peaking in the 66-72 hour window (Figure 4C and 4E and Movie S1). These transfers were accomplished as follows (Movie S1): the infected resident macrophage was approached by an uninfected peripheral monocyte. The cells then converged for a period of time before separating again, with the bacteria now being associated with the peripheral monocyte. Transfer events were not observed for PGL-deficient infection in the 66-72 hour window. (Figure 4E and Table S2). Thus, PGL-deficient bacteria largely remained within resident macrophages longer than wildtype bacteria. Furthermore, we documented clearance events of PGL-deficient *M. marinum* by the initially infected macrophage (Figure 4D and Movie S2). In contrast, clearance events were not observed during wildtype *M. marinum* infection.

To rigorously examine the kinetics of clearance in relation to the bacterial transfer events we had observed, we monitored ~ 30 animals for bacterial clearance by imaging them once every 24 hours. Since wildtype bacteria only transfer into permissive monocytes starting at 54 hours, the differential clearance of wildtype and PGL-deficient bacteria should become apparent only after this time-point. This was the case (Figure 4F). In contrast, PDIM-deficient bacteria started to be cleared within 24 hours (Figure 4F) consistent with their recruiting microbicidal monocytes within two hours and being phagocytosed by them within twelve hours (Figure 4A and 4B).

Imaging of these early mycobacterium-beneficial transfer events revealed they were distinct in their cellular morphology from subsequent intercellular bacterial transfer observed in the forming granuloma, which is dependent on the apoptotic death of the infected macrophage, the bacterial contents of which are engulfed by newly arriving macrophages (Davis and Ramakrishnan, 2009). In contrast the PGL-dependent transfer event was characterized by movement of the “donor” infected resident macrophage until the time that it converged with the “recipient” peripheral monocyte (Movie S1). Because the ESX-1 locus promotes apoptosis of infected macrophages (Davis and Ramakrishnan, 2009), our finding that ESX-1-deficient *M. marinum* were not compromised during early infectivity (Figure 3H), suggested that efferocytosis is not mediating this transfer event. To confirm this, we used the pan-caspase inhibitor QVD-OPH that reduces apoptotic cells substantially (7.2-fold) in the context of *M. marinum* infection of the zebrafish (Yang et al., 2012). QVD-OPH treatment did not reduce the early infectivity of *M. marinum* (Figure 4G), further suggesting that this transfer event is not dependent on efferocytosis. Rather, transfer was occurring between living cells, similar to the findings that, following intimate contact between macrophages in culture, intracellular Gram-negative pathogens can transfer between the two cells in a process known as trogocytosis (Steele et al., 2016).

Together, these findings are consistent with the model that PGL-competent bacteria transfer into the permissive monocytes they recruit. Conversely, our finding that PGL-deficient mycobacteria have a more prolonged sojourn in resident macrophages in which they are cleared, suggests that resident macrophages are more microbicidal than CCR2-recruited monocytes.

### Resident macrophages are more microbicidal than monocytes

Our findings linking increased time in the resident macrophage to increased bacterial killing suggested that resident macrophages are more microbicidal than CCR2- recruited monocytes. To address this question, we took advantage of our finding that following infection with 1-3 bacteria, only resident macrophages harbor PGL-deficient bacteria for at least the first 4.5 days (Figure 4E). We found that in PU.1 morphant animals lacking myeloid cells and therefore the resident macrophage niche they occupied at this stage, PGL-deficient bacteria were able to establish infection at wildtype levels (Figure 5A). These data further suggested that resident macrophages are microbicidal to PGL-deficient bacteria. We found that PGL-deficient infection resulted in more inducible nitric oxide synthase (iNOS)-positive cells than wildtype infection (Figure 5B). This was similar to the case of the PDIM-deficient mutant whose TLR-recruited monocytes express more iNOS than CCL2-elicited monocytes (Figure 5B) (Cambier et al., 2014b). However, since PGL-deficient *M. marinum* recruits only resident macrophages, the increased iNOS production must be coming from the resident macrophages - i.e. resident macrophages, like TLR-recruited monocytes, also produce more iNOS than CCL2- elicited permissive monocytes following infection. If this were the case then following delivery directly to monocytes via caudal vein infection (Figure 1A), PGL-deficient bacteria should result in the same low number of infected iNOS-positive cells as wildtype bacteria, and they did (Figure 5B). PDIM-deficient infection induced more iNOS in the caudal vein also (Figure 5B), suggesting that myeloid cells responding to PDIM-deficient bacteria are more activated regardless of location. Finally, we showed that the increased iNOS expression in the resident macrophages contributed to their increased microbicidal activity, as it does for TLR-recruited monocytes (Cambier et al., 2014b) - treatment of animals with the nitric oxide scavenger CPTIO increased the infectivity of PGL-deficient bacteria delivered into the HBV (Figure 5C). Together these results suggested that the reduced infectivity of PGL-deficient bacteria is due to their prolonged sojourn in resident macrophages. If so, then the infectivity of PGL-deficient bacteria should be restored when delivered directly to monocytes by intravenous infection. It was (Figure 5D), and this result further showed that mycobacterial PGL does not protect mycobacteria from the microbicidal activity of resident macrophages but rather promotes their escape into the more permissive monocytes. Both CCR2-deficiency and STING-deficiency, which produced the expected decreased in infectivity of wildtype *M. marinum* upon hindbrain ventricle infection, failed to do so when the bacteria were delivered directly to monocytes through caudal vein infection (Figure 5E and 5F). Together, these findings highlighted the role of STING and CCL2 as early host susceptibility factors that work by enabling recruitment of peripheral monocytes to sites of infection.

**Figure 5:**
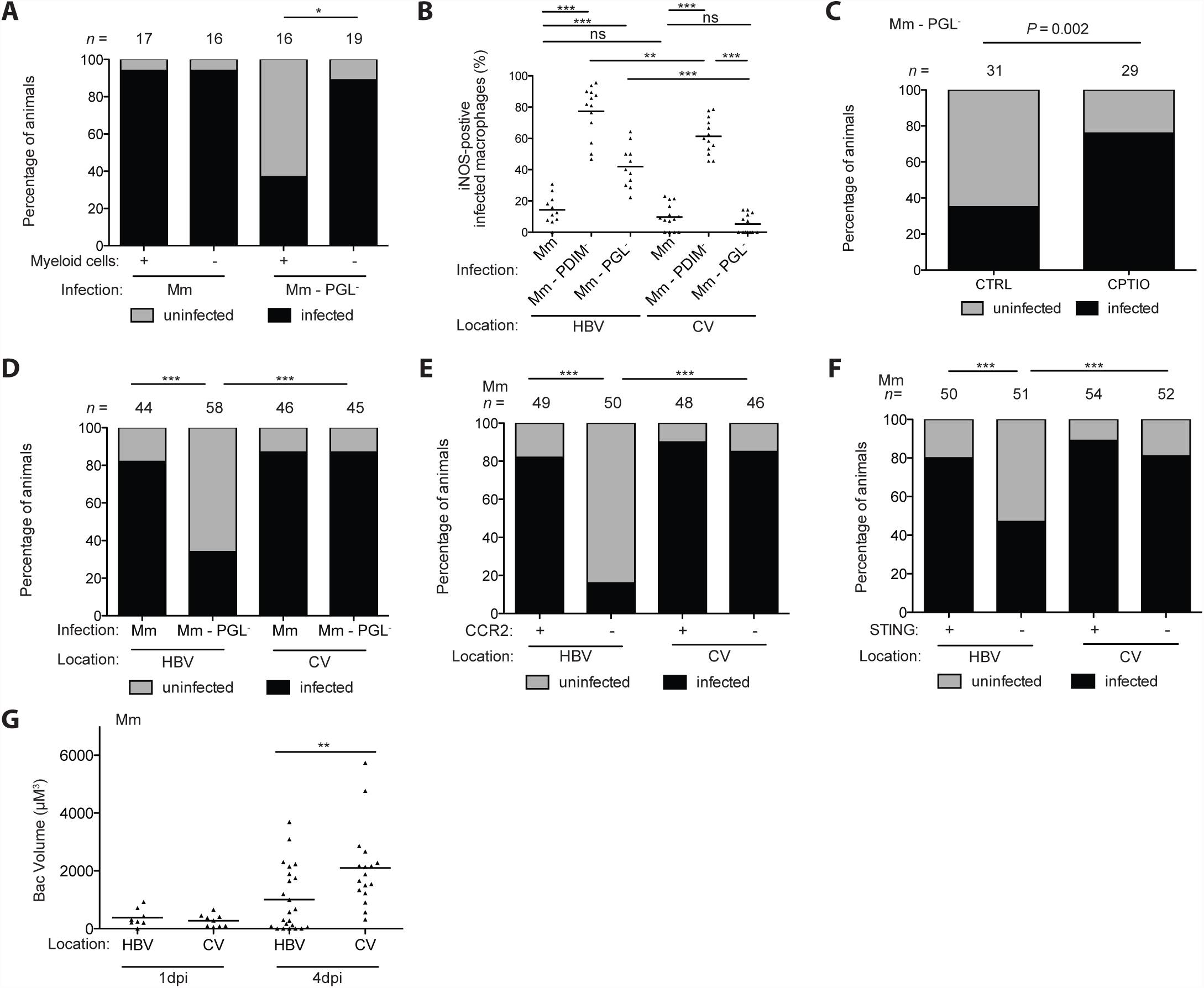
Resident macrophages are more microbicidal than monocytes. (A) Percentage of infected (black) or uninfected (gray) wildtype or myeloid-deficient fish at 5 dpi after HBV infection with 1-3 wildtype or PGL^-^ *M. marinum*. *n* = number of larvae per group. (B) Percentage of iNOS-positive infected myeloid cells in the HBV or CV at 3dpi with 80 wildtype, PDIM^-^ or PGL^-^ *M. marinum*. (C) Percentage of infected (black) or uninfected (gray) wildtype fish at 5 dpi after HBV infection with 1-3 PGL^-^ *M. marinum*. Control, CTRL; RNS scavenger (CPTIO), CPTIO. *n* = number of larvae per group. (D) Percentage of infected (black) or uninfected (gray) wildtype fish at 5 dpi with 1-3 wildtype or PGL^-^ *M. marinum* into the HBV or CV. *n* = number of larvae per group. (E) Percentage of infected (black) or uninfected (grey) wildtype or CCR2-deficient fish at 5 dpi with 1-3 wildtype *M. marinum* in the HBV or CV. *n* = number of larvae per group. (F) Percentage of infected (black) or uninfected (grey) wildtype or STING-deficient fish at 5 dpi with 1-3 wildtype *M. marinum* in the HBV or CV. *n* = number of larvae per group. (G) Mean bacterial volume at 1 and 4dpi with a single wildtype *M. marinum* bacterium in the HBV or CV of wildtype fish. Results in (A)-(G) representative of three experiments. (B) and (G) significance testing done using one-way ANOVA, with Bonferroni’s post-test for comparisons shown. (A) and (C) – (F) significance testing done using Fisher’s exact test for the comparisons shown. \**P* < 0.05, \*\**P* < 0.01, \*\*\**P* < 0.001.

Finally, we asked if the 54-90 hour sojourn in resident macrophages was at all detrimental to wildtype bacteria. The infectivity assay we had used so far only assessed whether the animals had cleared the bacteria or not, and not the extent of bacterial growth in the animals that did not clear them. We now tested this following infection of animals with a single bacterium. We found twice as much bacterial growth in the caudal vein compared to the HBV (Figure 5G). Together these results show that resident macrophages are more microbicidal than the permissive monocytes to which the wildtype bacteria eventually gain access. Moreover, the resident macrophage plays a growth-restrictive role even to wildtype PGL-expressing bacteria during the truncated time period that they remain in it.

### Human alveolar macrophages rapidly secrete CCL2 after mycobacterial infection in a PGL-dependent fashion

Our prior work had shown that pathogenic mycobacteria establish infection by recruiting and infecting permissive monocytes while having specialized strategies to avoid recruiting microbicidal cells - neutrophils (Yang et al., 2012), and TLR-stimulated monocytes (Cambier et al., 2014b). The latter strategy requires that mycobacteria initiate infection in the lower lung, so as to avoid the TLR-stimulated microbicidal monocytes by the mucosal flora of the upper airway. The present work had now identified the resident macrophage as another default rapid first-responder microbicidal cell that mycobacteria cannot avoid even in the lower airways, and must therefore co-opt them into their escape strategy by inducing them to secrete CCL2 (Figure 6). In terms of human relevance of our zebrafish findings, our findings that resident macrophages are more microbicidal than peripheral monocytes already had support from human studies: human alveolar macrophages have substantial mycobactericidal activity ex vivo, in contrast to peripheral blood monocytes which not only fail to kill mycobacteria but are growth-permissive (Aston et al., 1998; Hirsch et al., 1994; Rich et al., 1997; van Zyl-Smit et al., 2014). Moreover, consistent with our findings, the microbicidal activity of human alveolar macrophages is at least in part mediated by nitric oxide (Hirsch et al., 1994).

**Figure 6:**
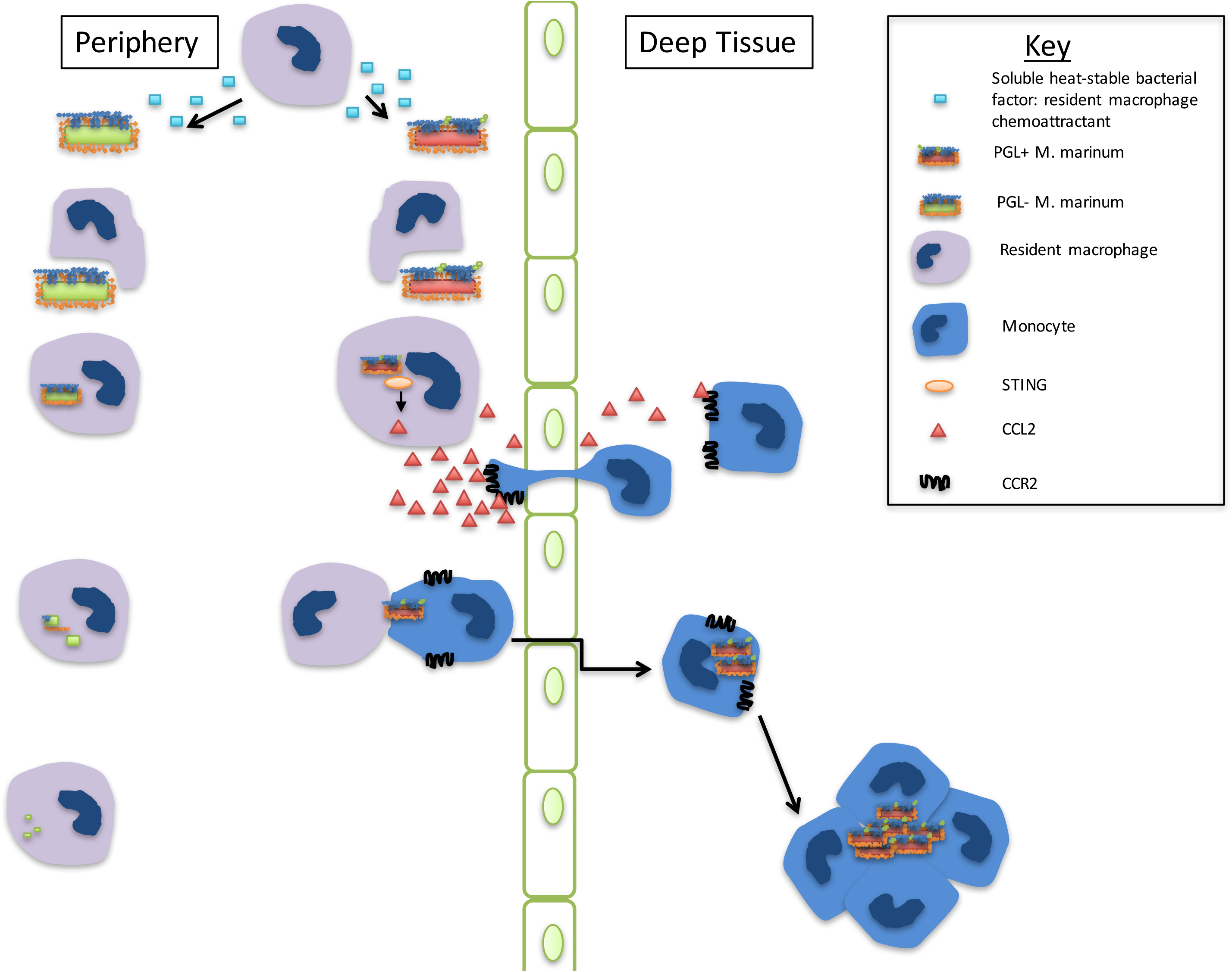
Schematic model of resident macrophage recruitment and subsequent CCL2 induction by mycobacterial PGL. Resident macrophages are the first cell type to respond to bacterial infections including mycobacteria, by responding to a soluble secreted bacterial factor. Mycobacterial PGL engages the cytosolic host protein STING to drive resident macrophages to make CCL2. This in turn recruits bacterium-permissive CCR2-positive monocytes, into which the mycobacteria transfer, in order to establish infection. In the absence of PGL, mycobacteria are cleared more often by resident macrophages.

Our model would further predict that human alveolar macrophages would rapidly produce CCL2 upon mycobacterial infection in a PGL-dependent fashion. To test this prediction, we performed a pilot experiment with human alveolar macrophages obtained by bronchoalveolar lavage. We infected them with either PGL-expressing or PGL-deficient *M. marinum*. CCL2 was induced in a PGL-dependent fashion at 60 minutes post-infection (Figure 7A and 7B - Donor 320). We then recruited 12 additional donors and infected their alveolar macrophages with PGL-expressing or PGL-deficient mycobacteria as well as with LPS (100ng/ml), a known CCL2 inducer. LPS induced CCL2 (>1.2 fold over uninfected) in 5 of 12 donors suggesting that the remaining were not capable of inducing CCL2 rapidly in response to a known inducer (Table S3). The LPS-nonresponding macrophages also did not induce CCL2 upon mycobacterial infection (Table S3). This nonresponsiveness is consistent with significant donor variation in human alveolar macrophage chemokine/cytokine secretion after mycobacterial infection (Keane et al., 2000). Of the LPS-responding macrophages, four of five induced CCL2 upon mycobacterial infection and this response was PGL-dependent (Figure 7A and 7B, and Table S3). In order to see if CCL2 induction occurred even earlier than 60 minutes, we had collected supernatants at 30 minutes. Only those donor alveolar macrophages that induced CCL2 in response to LPS and mycobacterial infection at the 60 minute time point, did so at the 30 minute time point (Figure 7C and Table S3). Again, CCL2 induction was PGL-dependent (Figure 7C and 7D). These experiments suggest that the rapid induction of CCL2 in human alveolar macrophages in response to mycobacterial infection is PGL-dependent.

**Figure 7:**
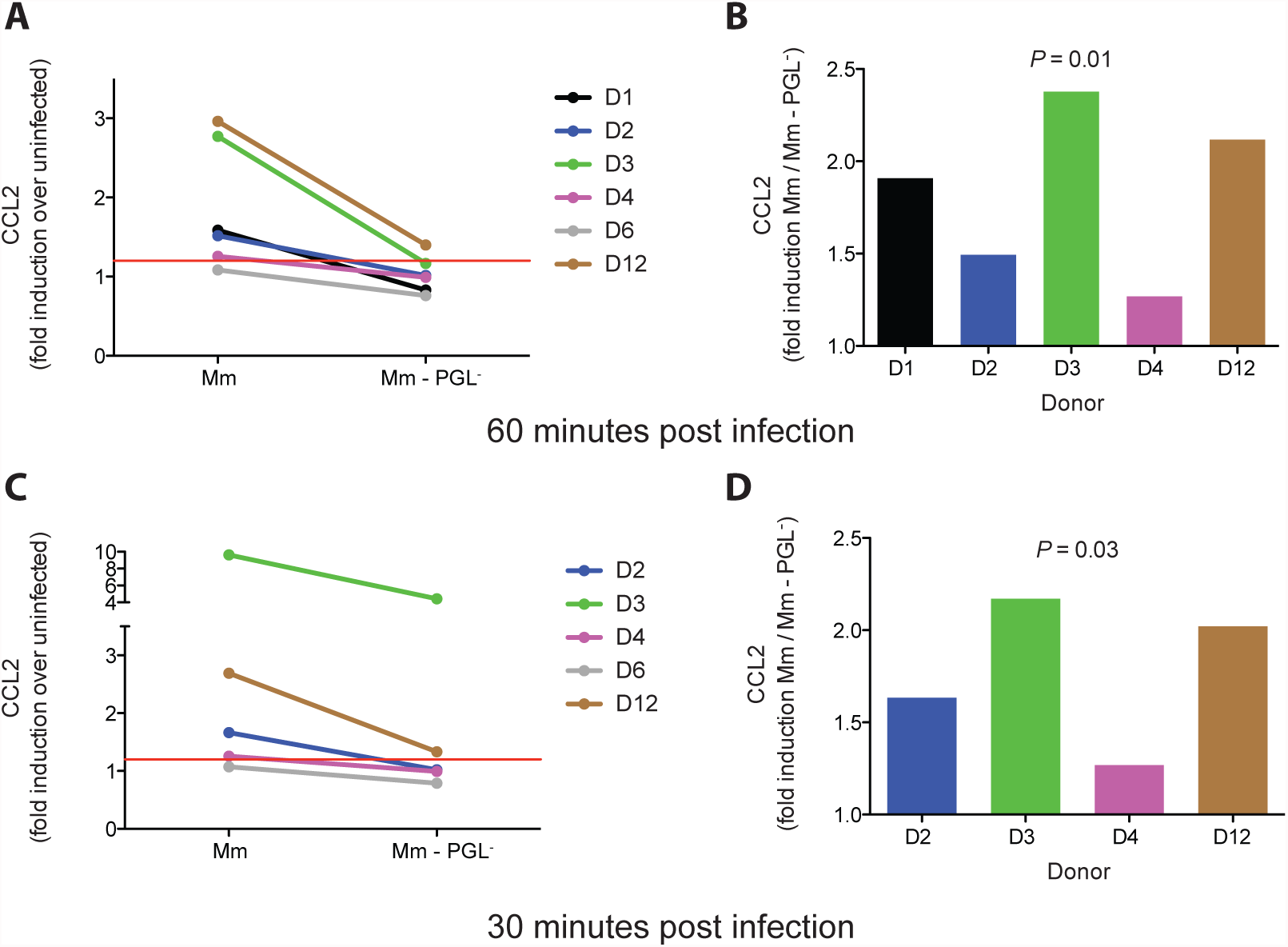
PGL-dependent CCL2 protein production following *M. marinum* infection of human alveolar macrophages. (A and C) Fold increase (over uninfected cells) in CCL2 protein levels in the supernatant of primary human alveolar macrophages following a 60-minute (A) or 30-minute (C) infection with wildtype *M. marinum*) or PGL-deficient *M. marinum*. (B and D) The same data as in (A and C) analyzed as fold increase in CCL2 of wildtype *M. marinum* over PGL- *M. marinum* at 60-minutes (B) and 30-minutes (D) post infection. Significance testing done using a one sample *t*-test to a hypothetical value of 1, corresponding to the null hypothesis that PGL does not influence CCL2 production following infection.

## DISCUSSION

By tracking the dynamics and kinetics of the earliest myeloid cell responses in the first hours of mycobacterial infection, we find that tissue resident macrophages are the first cells to come in contact with any infecting bacteria in response to a ubiquitous heat-stable secreted bacterial signal (Figure 6). Arriving to virulent mycobacteria, they rapidly infect them and are capable of eradicating them (Figure 6). In turn, mycobacterium’s counterstrategy to circumvent this first-line host defense that it cannot evade is to engineer its escape from these cells (Figure 6).

From a teleological perspective, these findings now explain why mycobacteria must deploy two distinct mechanisms for macrophage recruitment - PGL-CCL2- mediated initially followed by ESX-1-MMP9-mediated in the forming granuloma (Volkman et al., 2010). Intercellular bacterial transfer in the granuloma requires the apoptotic death of a highly-infected macrophage that is then engulfed by multiple new recruits so as to expand the bacterial niche (Davis and Ramakrishnan, 2009). Therefore, this mechanism of granuloma expansion depends upon bacteria being in a growth-permissive cell. Our new findings show that PGL-induced CCL2 occurs even under the bacteriostatic or bactericidal conditions imposed by the resident macrophage, allowing even the few remaining bacteria to escape into permissive cells. We have recently shown that *Mycobacterium leprae*’s PGL-1, differing from *M. marinum*’s and *M. tuberculosis’* PGL in the carbohydrate domain, is required for monocyte-mediated demyelination at a later step of the infection (Madigan et al, in revision). However, *M. leprae* also mediates recruitment of monocytes through CCL2/CCR2 signaling, suggesting that its specialized PGL-1 still retains the basal function of eliciting permissive monocytes to promote its infectivity at the first steps of infection. It is interesting that both PGL-mediated functions - establishment of infection and demyelination - are through manipulation of host myeloid cells (Madigan et al, in revision, and this work).

Our findings highlight not only both phylogenetic and ontogenic conservation of resident macrophage function but also suggest that different tissue resident macrophages - even the most specialized brain resident macrophages (Casano and Peri, 2015) - all retain their primal function as sentries against invading pathogens (Epelman et al., 2014; Gordon et al., 2014). The finding that resident macrophages can make short shrift of mycobacteria, notoriously pernicious pathogens, is particularly noteworthy given their key role in tissue homeostasis (Epelman et al., 2014). Then again, it is curious that CCL2-elicited monocytes provide a safe-haven to mycobacteria as CCR2^+^ monocytes are broadly microbicidal against bacterial, fungal protozoan and viral pathogens (Serbina et al., 2008). Indeed, these cells, also called inflammatory monocytes, are implicated in the pathogenesis of multiple inflammatory diseases affecting the brain, gut and vascular system (Lauvau et al., 2014; Shi and Pamer, 2011). On the other hand, CCR2^+^ myeloid cells have been implicated in promoting an immunosuppressive tumor environment (Lesokhin et al., 2012). Our data identify a permissive role for these cells in the context of an important intracellular infection. Consistent with our findings, CCL2-recruited monocytes have been previously shown to be more permissive to *M. tuberculosis* growth in the lungs of mice (Antonelli et al., 2010), and mice overexpressing CCL2 were found to be more susceptible to challenge with *M. tuberculosis* (Rutledge et al., 1995). Their reduced microbicidal capacity in response to mycobacterial infection may simply reflect the masking of activating TLR ligands by mycobacteria, though it is notable that even in the absence of TLR-mediated activation, resident macrophages are more microbicidal to mycobacteria than monocytes. Of course TB is a complex infection and it is possible that as infection progresses, these same inflammatory monocytes could take on a host-beneficial role in delivering mycobacterial antigens to pulmonary lymph nodes to eventually lead to antigen specific T-cell responses (Samstein et al., 2013). However, even this role may have complex consequences - while T cell responses are clearly protective for individuals, they may also be paradoxically benefitting bacteria by promoting transmission to new individuals (Comas et al., 2010). Overall, our findings add to the discussion of the plasticity and context-dependent function of myeloid cells, for which there is increasing appreciation particularly with the advent of in vivo studies suggesting that myeloid cell functions defy rigid classifications (Martinez and Gordon, 2014; Murray et al., 2014).

Finally, we note that while evolutionary ancestors of *M. tuberculosis* e.g. *M. marinum* and *Mycobacterium cannetti* uniformly express PGL, the prevalence of PGL-expression in modern-day *M. tuberculosis* strains is not clear (Gagneux, 2006; Pang et al., 2012). This work emphasizes the need to assess the prevalence of PGL-positive strains, and to thoroughly examine TB transmission epidemiology in regions where PGL-expressing strains abound, while devising therapeutic strategies to block PGL to prevent TB infection and transmission.

## STAR METHODS

### Bacterial Strains and Methods

*M. marinum* strain M (ATCC BAA-535) Δ*mmpL7,* Δ*pks15,* and Δ*esx-1* mutants expressing either TdTomato or Wasabi under the control of the *msp12* promoter (Cambier et al., 2014b; Takaki et al., 2013) were grown under hygromycin (Mediatech) selection in 7H9 Middlebrook’s medium (Difco) supplemented with oleic acid, albumin, dextrose, and Tween-80 (Sigma). To prepare heat-killed *M. marinum,* bacteria were incubated at 80**°**C for 20 minutes. To prepare bacterial supernatants, bacteria were grown to an OD600 of 0.6, pelleted and the supernatant was then filtered twice through a 0.2μm filter. The *P. aeruginosa* PAO1 fluorescent strain has been described (Brannon et al., 2009). The *S. aureus* Newman strain expressing pOS1-SdrC-mCherry #391 was a gift from Dr. Juliane Bubeck Wardenburg.

### Bead Injections

Sterile red-fluorescent 1μm beads (Thermo-Fisher Scientific F8821) were diluted ten fold with sterile PBS resulting in 3.64 x 10^3^ beads/nL. Approximately 5 nL of the bead mixture was injected into the hindbrain ventricle of 2 dpf larvae for a total of 1.8 x10^4^ beads per larva.

### QVD-OPH and CPTIO Treatment

CPTIO or QVD-OPH (Sigma) was used at a final concentration of 50 mM and 50 μM, respectively, in 0.5% dimethylsulphoxide in fish water. Fish were incubated immediately following infection and fresh inhibitor was added every 24 h until experiment end point.

### Zebrafish Husbandry and Infections

Wildtype AB and *csf1ra^j4blue^* homozygous mutant (*csf1r*^-/-^) zebrafish (Parichy et al., 2000) were maintained as described (Takaki et al., 2013). The Tg(*mpeg1:YFP)^w200^*, and Tg(*mpeg1:Brainbow)^w201^* (expressing tdTomato) lines were used as previously described(Pagán et al., 2015)Larvae (of undetermined sex given the early developmental stages used) were infected at 48 hours post-fertilization (hpf) via caudal vein or hindbrain ventricle injection using single-cell suspensions of known titer (Takaki et al., 2013; 2012). Number of animals to be used for each experiment was guided by pilot experiments or by past results with other bacterial mutants and/or zebrafish. Larvae were randomly allotted to the different experimental conditions. All experiments where *csf1r*^-/-^ zebrafish were used, *csf1r*^-/-^ were either in-crossed or outcrossed to wildtype ABs to generate *csf1r^+/-^* which are phenotypically wildtype (Pagán et al., 2015). Zebrafish husbandry and all experiments performed on them were in compliance with guidelines from the UK Home Office (Cambridge experiments) and in compliance with the U.S. National Institutes of Health guidelines and approved by the University of Washington Institutional Animal Care and Use Committee (Seattle experiments) and the Stanford Institutional Animal Care and Use Committee (Stanford experiments).

### Confocal Microscopy and Image-Based Quantification of Infection

Larvae were embedded in 1.5% agarose (low melting point) (Davis and Ramakrishnan, 2009). A series of z-stack images with a 2 μm step size was generated through the infected HBV, using the galvo scanner (laser scanner) of the Nikon A1 confocal microscope with a 20x Plan Apo 0.75 NA objective. Bacterial burdens were determined by using the 3D surface-rendering feature of Imaris (Bitplane Scientific Software) (Yang et al., 2012).

### Hindbrain Kinetic Assays

Macrophage recruitment assays were performed as previously described (Takaki et al., 2012). For assays distinguishing resident macrophages from monocytes, 200 μg/ml Hoechst 33342 (Invitrogen) was injected via the caudal vein as previously described (Davis and Ramakrishnan, 2009) 2 hours prior to infection into the hindbrain. Differential interference contrast and fluorescent imaging using Nikon’s Eclipse E600 was done every ~30 min to identify resident macrophages (Hoechst/blue fluorescence negative) and monocytes (Hoechst/blue fluorescence positive). Objectives used in this assay included 20x Plan Fluor 0.5 NA and 40x Plan Fluor 0.75 NA.

### Morpholinos

The STING morpholino 5’TGGAATGGGATCAATCTTACCAGCA3’ was designed to block the exon 2 intron 2 border. The following primer pair 5’CTGCTGGACTGGGTTTTCTTACTC3’ and 5’TGGGTGATCTTGTAGACGCTGTTA3’ was used to assess morpholino efficiency. STING morpholino injection led to nonsense mediated decay of mRNA transcripts out to 5dpf. The STING morpholino and the CCR2, PU.1, and MyD88 morpholinos previously described (Cambier et al., 2014b) were injected into the 1-4 cell stage of the developing embryo (Tobin et al., 2010).

### Quantitative Real-time PCR (qRT-PCR)

Total RNA was isolated from pools of 20-40 larvae as previously described (Clay et al., 2007), using TRIzol Reagent (Life Technologies) and was used to synthesize cDNA with Superscript III reverse transcriptase and oligo DT primers (ThermoFisher Scientific). Quantification of *ccl2* RNA levels were determined using SYBR green PCR Master Mix (Applied Biosystems) on an ABI Prism 7300 Real-Time PCR System (Applied Biosystems) using the following primer pair; 5’GTCTGGTGCTCTTCGCTTTC3’ and 5’TGCAGAGAAGATGCGTCGTA3’. Average values of technical triplicates of each biological replicate were plotted. Data were normalized to *β-actin* for ΔΔCt analysis.

### Infectivity Assay

2 dpf larvae were infected via the hindbrain ventricle with an average of 0.8 bacteria per injection as previously described (Cambier et al., 2014b). Fish harboring 1-3 bacteria for some experiments or 1 bacterium for others were identified at 5 hours post infection by confocal microscopy. These infected fish were then evaluated at 5 dpi, or every 24 hours following infection, and were scored as infected or uninfected, based on the presence or absence of fluorescent bacteria.

### CCL2 In Situ Hybridization

In situ hybridization was performed as previously described(Clay et al., 2007). Zebrafish CCL2 (ENSDARG00000041835) was cloned from adult pooled cDNA constructed from isolating RNA from homogenized adult tissues using Trizol (ThermoFisher), chloroform extraction and purification using RNeasy mini kit (Qiagen). Superscript III reverse transcriptase (ThermoFisher) was used to make cDNA and the following primer pair 5’GTCAGCTAGGATCCATGAGGCCGTCCTGCATC C3’ and 5’GTCAGCTATCTAGATTAGGCGCTGTCACCAGAG3’ was used to clone zebrafish CCL2.

### Human Alveolar Macrophage Collection

Human alveolar macrophages (AMs) were retrieved at bronchoscopy as approved by the Research Ethics Committee of St. James’s Hospital, and previously reported(Berg et al., 2016; O’Leary et al., 2014). Briefly all donors were patients undergoing clinically indicated bronchoscopy and written informed consent for retrieving additional bronchial washings for research was obtained prior to the procedure. Bronchial washing fluid was filtered through a 100μm nylon strainer (BD Falcon, BD Bioscience, Belgium) and centrifuged at 390g for 10min. Alveolar macrophages were resuspended in RPMI 1640 culture media supplemented with 10% fetal bovine serum (FBS, Gibco), 2.5ug/ml fungizone and 50 μg/ml cefotaxime. AMs were seeded at a density of 5 x 10^4^ cells/well in 96-well plates (Corning Costar ^TM^, Nijmegen, Netherlands). AMs were purified by plastic adherence, non-adherent cells were removed by washing after 24hrs.

### Infection of Human Alveolar Macrophages

On the day of infection *M. marinum* wild-type and Δ*pks15* growing in Middlebrook 7H9 medium were centrifuged at 2900g for 10min and resuspended in RPMI 1640 containing 10% FCS. Clumps were disrupted by passing the bacilli through a 25-gauge needle 6-8 times and the sample was centrifuged at 100 (x)g for 3 min to remove any remaining clumps. To assess the adequacy of dispersion and to determine the MOI, macrophages were infected with varying amounts of resuspended *M. marinum* wild type and PGL-deficient for 2hrs. Extracellular bacteria were washed off, and cells were fixed with 2% paraformaldehyde for 10mins. Macrophage nuclei were counterstained with 10μ/ml of Hoechst 33358 (Sigma). The percentage of infected cells and the number of bacilli per cell were determined by fluorescent microscopy (Olympus IX51, Olympus Europa GmbH, Germany) for each donor, as previously described(Gleeson et al., 2016; O’Leary et al., 2011; O’Sullivan et al., 2007; O’Leary et al., 2014). Based on this result alveolar macrophages were infected at an estimated MOI of 1-10 bacilli. At 1hr post-infection supernatants were harvested for MCP-1 assay.

### MesoScale Discovery Chemokine (CCL2 (MCP1)) Assay

Human MCP-1 chemokine kit (Meso Scale Discovery®, Maryland, USA) was used as per manufacturers’ instructions, briefly samples, standards and controls were added at 25 μL per well. Detection antibody was added at 25 μL per well, 150 μL of the MSD Read Buffer was added to each well and the MSD plates were measured on the MSD Sector Imager 2400 plate reader. The raw data was measured as electrochemiluminescence signal (light) detected by photodetectors and analyzed using the Discovery Workbench 3.0 software (MSD). A 4-parameter logistic fit curve was generated for CCL2/MCP1 using the standards and the concentration of each sample calculated.

### Statistics

Statistical analyses were performed using Prism 5.01 (GraphPad). Error bars represent standard error of mean. Post-test *P* values are as follows: \**P* < 0.05; \*\**P* < 0.01; \*\*\**P* < 0.001

## AUTHOR CONTRIBUTIONS

C.J.C. and L.R. conceived, designed and analyzed the zebrafish experiments and CJC performed them. C.J.C, S.M.O., M.P.O., J.K., L.R. designed and analyzed the human experiments, and S.M.O performed them. C.J.C. and L.R. wrote the paper. All authors edited the paper.

## ACKNOWLEDGMENTS

We thank C.R. Bertozzi for providing space and resources to complete this project, D. Stetson for suggesting STING, A. Pagán for suggesting the human experiments to test the model, K. Urdahl for discussion and suggestions, S. Candel, J.M. Davis. P. Edelstein, S. Falkow, D. Tobin and K. Urdahl for manuscript review, and J. Cameron, R. Keeble and N. Goodwin for zebrafish husbandry. For the human work, we thank Drs. F. O’Connell, AM McLaughlin, the research nurses of the Wellcome Trust-HRB Clinical Research Facility, and the staff and patients of the St. James’s Hospital Bronchoscopy Clinic, Dublin, and Karl Gogan for assistance in preparation of alveolar macrophages.

This work was supported by NIH grant R37AI054503 and the National Institute of Health Research Cambridge Biomedical Research Centre (L.R), and NIH training grant T32 AI55396 (C.J.C.), the Health Research Board of Ireland (S.O’L., M.P.O’S. and J.K.), and The Royal City of Dublin□Hospital Trust (J.K.). L.R. is a Wellcome Trust Principal Research Fellow and C.J.C. is a Damon Runyon Postdoctoral Fellow.

---

**Movie S1:** Transfer of green Wasabi-expressing wildtype Mm from red Mpeg1+ resident macrophage to blue Hoechst+ red Mpeg1+ monocyte. Imaged every 10min.

**Movie S2:** Green Mpeg1+ cell infected with red TdTomato-expressing PGL - Mm. Surface rendering done by setting a threshold for the red fluorescence using Imaris. Imaged every 10min.

**Table S1:** The number of resident macrophages and monocytes responding to infection with 1-3 wildtype Mm or PDIM- Mm every 10min from 1 – 11 hours post infection. Raw data used to make Figure 4A.

**Table S2:** Number of transfer events occurring during the first 4.5 days following infection with 1-3 bacteria. The number of fish imaged, fish remaining infected, number of total infected macrophages, and the number of transfer events recorded during each time-period of imaging are provided following infection with either wildtype or PGL^-^ *M. marinum* (Mm). Raw data for Figure 4E.

**Table S3**: CCL2 production by human alveolar macorphages. A complete list of donor alveolar macrophages and their CCL2 production in response to LPS, wildtype Mm, and PGL- Mm.

